# Transcriptome analysis reveals high tumor heterogeneity with respect to re-activation of stemness and proliferation programs

**DOI:** 10.1101/2021.09.02.458724

**Authors:** Artem Baranovsky, Timofei Ivanov, Marina Granovskaya, Dmitri Papatsenko, Dmitri Pervouchine

## Abstract

Significant alterations in signaling pathways and transcriptional regulatory programs together represent major hallmarks of many cancers. These, among all, include the reactivation of stemness, which is registered by the expression of pathways that are active in the embryonic stem cells (ESCs). Here, we assembled gene sets that reflect the stemness and proliferation signatures and used them to analyze a large panel of RNA-seq data from The Cancer Genome Atlas (TCGA) Consortium in order to specifically assess the expression of stemness-related and proliferation-related genes across a collection of different tumor types. We introduced a metric that captures the collective similarity of the expression profile of a tumor to that of ESCs, which showed that stemness and proliferation signatures vary greatly between different tumor types. We also observed a high degree of intertumoral heterogeneity in the expression of stemness- and proliferation-related genes, which was associated with increased hazard ratios in a fraction of tumors and mirrored by high intratumoral heterogeneity across cancer cells in single cell RNA-seq datasets. Taken together, these results indicate that the expression of stemness signatures is highly heterogeneous and cannot be used as a universal determinant of cancer. This calls into question the universal validity of diagnostic tests that are based on stem cell markers.

**Author summary:** Cancer is a deadly human disease which is characterized by uncontrolled proliferation of abnormal cells. Stemness, or the expression of stem cell markers, has been identified as a key feature for cancer progression and in many cases correlates with patient survival. In this work, we reanalyzed a large cohort of cancers from TCGA (The Cancer Genome Atlas) collection and single-cell cancer data, and found that the degree of stemness reactivation varies greatly between different tumor types, between different tumors of the same type, and also different cells within a tumor. The observed stemness heterogeneity implies that the expression of stemness markers cannot be used as a universal determinant of cancer and calls into question the validity of stemness-based tests that are frequently used for cancer diagnostics.

## Introduction

Cancer is one of the major causes of death worldwide [1]. A significant progress in cancer treatment has been achieved with the development of a large therapeutic arsenal including chemotherapy, surgery, radiation therapy, and immunotherapy [2–4]. An important direction in clinical research focuses on the so-called Cancer Stem Cells (CSC), a subpopulation of tumorous cells with high tumor-initiating potential [5]. CSCs are increasingly regarded as prominent targets for anti-cancer therapy, although the degree of expression of the stem cell-like phenotype, referred to as stemness, may vary between different tumors [6].

Several hypotheses relate stemness with the origin of cancer. It is acknowledged that cancers arise either from a malignant transformation of a progenitor cell, or from a non-stem cell, which reacquired the stemness potential [7–9]. This paradigm is sustained by significant convergence of stem cells (SC) and CSCs in the activated signaling cascades, moreover in their overlapping expression of a set of biomarkers encompassing the classical self-renewal-associated pathways Wnt/*β*-catenin, Bmi-1, sonic hedgehog, Notch, and PTEN [10]. Additionally, both SCs and CSCs express tissue-specific stem cell markers [11–14]. Such concordant molecular profile stipulates key aspects of SC and CSC phenotype including longevity, dormancy, niche dependence, and the potential for asymmetric cell division [15–18].

Tumors display frequent inter- and intratumor heterogeneity in alterations of the global gene expression programs and, in particular, in the expression of stemness markers [19–22]. Commonly used CSC-associated markers have a variable expression in glioblastoma [23]. The expression of breast cancer stem cell markers, ALDH1A1 and CD133, shows spatial heterogeneity across patients [19]. Stemness phenotype is also heterogeneous among many normal adult SC populations in the human body, where the SCs uphold tissue regenerative capacity [24, 25]. Moreover, the degree of activation of stemness programs is associated with increased expression of multiple immunosuppressive pathways and decreased anticancer immunity [26]. This suggests that the stemness state may itself be highly heterogeneous within and between tumor types, which may play an important role in cancer pathogenesis.

A number of metrics have been developed to quantify stemness [20, 26, 27]. These metrics correlate with intratumoral heterogeneity, antitumor immune response, and clinical prognosis [20, 26, 27]. However, the heterogeneity of the stemness signature has not been assessed in detail [19]. Here, we reanalyzed the transcriptomes of 19 tumor types from The Cancer Genome Atlas (TCGA) Consortium and juxtaposed them with the transcriptomes of human embryonic stem cells [28], adult stem cells [29, 30], and induced pluripotent stem cells (iPSC) within 4 days of differentiation [31]. The comparison of these transcriptomes from the perspective of reactivation of the stemness program revealed a pronounced heterogeneity both within and between tumor types, which in many cases was associated with tumor-specific patient survival. To interrogate intratumoral heterogeneity, we additionally demonstrated an increased variability of stemness signature in cancer cells compared to non-cancer cells using single cell transcriptomes of colorectal cancer [32].

## Materials and methods

### RNA-seq data and related clinical data

Gene expression values for tumors and normal tissues from the TCGA dataset and all relevant metadata were downloaded in the form of read counts using R package `TCGAbiolinks` [33] (S1 Data File). The fastq files for stem cell (SC) datasets comprising iPSC (fibroblasts purified from skin-punch biopsies from six males and six females that were reprogrammed using transfection with an episomal plasmids containing OCT3/4, SOX2, KLF4, L-MYC, LIN28, and an shRNA against p53 [34]), ESC, other types of pluripotent stem cells (PSC), and adult stem cells (ASC) were either downloaded from Sequence Read Archive (SRA) or obtained by direct download from arrayexpress ftp server (S1 Table). All fastq files were processed by a uniform pipeline, in which the reads were first checked for quality, trimmed using `TrimGalore`, and mapped with STAR version 2.7.1a [35] to the December 2013 assembly of the human genome (hg38, GRCh38). Gene expression levels were quantified using the Subread tool [36] implemented in `Rsubread` package version 1.34.7 [37]. The read count matrices from TCGA and stem cell datasets were merged, filtered by gene expression, and normalized using edgeR [38]. The resulting combined matrix was corrected for tumor purity as explained below. The raw UMI counts for the scRNA-seq dataset (63,689 cells from 23 primary colorectal cancer and 10 matched normal mucosa samples) were downloaded from GEO under the accession GSE132465 [32]. The counts were normalized for sequencing depth per cell and transformed to log_2_-scale using a pseudocount of one. The subsequent analysis and visualization of single cell RNA-seq data were done using R package `Seurat` version 4.0.2.

### Correction for tumor purity

To mitigate the influence of normal cells on gene expression in tumor samples, we regressed out the effect of tumor purity defined by consensus tumor purity estimate (CPE). First, we downloaded a table of precomputed CPE values which is available as a Supplementary Data 1 in [39]. Next, we filtered out tumor samples where CPE was not available and built a collection of linear models of the form *Y*_*i,s*_ *∼ β*_*i*,0_ + *β*_*i*,1_ *∗ P*_*s*_ + *ε*_*i,s*_ individually for each gene *i*, where *Y*_*i,s*_ is the TMM-normalized log_2_(*CPM*) of the gene *i* in the sample *s*, and *P*_*s*_ is the value of the CPE of the sample *s*, and *ε*_*i,s*_ is the error term. The purity-corrected TMM-normalized log_2_(*CPM*) of gene *i* in the sample *s* were obtained from *Y*_*i,s*_ by subtracting the linear part, i.e., 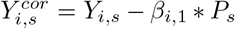 Since the CPE metric provides an estimate for the proportion of tumor cells, which are presumably absent in normal tissues, we assigned a random value close to zero from normal distribution with the mean 0.08 (0.05 percentile of CPE distribution) and standard deviation 0.03 (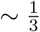 of the 0.05 percentile of CPE distribution) to CPE of all normal tissues and iPSCs. The resulting values of 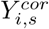 showed no statistically significant dependence on CPE (mean *R*^2^ *⋍* 0).

### Differential gene expression analysis

The `edgeR` package was used for differential expression analysis. Raw read counts were normalized by `edgeR` using the TMM (Trimmed Mean of M-values) method [38]. The model was fitted to the data using `glmQLFit` function. Differential gene expression was estimated with `glmTreat` function. For GO enrichment analysis, differentially expressed genes were defined by | log_2_(*FC*)| *≥* 1 and adjusted p-value cutoff of 0.05.

### Gene Ontology analysis

Gene Ontology analysis of differentially expressed genes was done using R package `clusterProfiler` [40] with the default parameters.

### Signature gene sets

The set of stemness gene signatures (*n* = 271) was originally designed by DPa by a combination of literature search and analysis of transcriptomic datasets. However, his efforts were interrupted by force majeure circumstances, and the rest of the team had to reassemble a similar set using another procedure outlined below.

We considered datasets spanning various experimental protocols, e.g. tissue-gene expression atlases, ESC differentiation time series, and knockdowns of pluripotency-associated transcription factors (ppTF-KDs) [41–44]. The datasets were analyzed one at a time, with individually selected cutoffs (S2 Table). The microarray gene expression data were downloaded using the R package `GEOquery` [45], normalized using the `rma` function available in the R package `affy` [46], log_2_-transformed and processed following the guidelines from the `affy` package vignette.

Our collection included two gene expression atlases, GSE1133 and GSE10246, both containing gene expression data from various mouse tissues and cell lines. We identified a group of samples, in which the stemness program could be active (S2 Table), and selected only one group in each atlas that was closest to ESCs (Blastocysts in GSE1133 and mouse ESCs in GSE10246). In each atlas, we computed log_2_ *FC* between the selected group and the rest of tissues or cell lines and computed *E*, the log-sum of gene expression values across samples. Then, we selected genes that satisfied the following conditions in at least *n*_*comp*_. of the comparisons in each atlas: log_2_ *FC >* 0.05, *E >* 3, *n*_*comp*_. = 34 for GSE1133, and log_2_ *FC >* 0.1, *E >* 4.5, *n*_*comp*_ = 49 for GSE10246. This resulted in two lists of putative stemness genes with 5764 and 2670 elements for GSE1133 and GSE10246, respectively.

Next, we analyzed a collection of ESC-line differentiation datasets, which included time series analysis of 14 days of differentiation in three mouse ESC-lines: J1, R1 and v6.5. As before, we computed *E* and log_2_ *FC* between the first (0 hours) and last (14 days) time points, separately for each cell line. The cutoffs log_2_ *FC >* 0.05 and *E >* 3 resulted in three lists of 3286, 3567, and 2609 putative stemness genes corresponding to J1, R1 and v6.5 lines, respectively. A similar analysis of shRNA knockdowns of five core pluripotency transcription factors, *POU5F1, NANOG, SOX2, ESRRB* and *SALL4*, in mouse ESCs resulted in 8588 genes with log_2_ *FC >* 0.05 and *E >* 3 in at least one of the experiments.

The intersection of these lists consisted of *n* = 454 genes, to which we added a set of manually collected and curated putative stemness genes, which were not picked by our analysis (*n* = 19) (S2 Data File). Then, we removed tissue-specific genes [47] (*n* = 7), proliferation signature genes (*n* = 49, see below), and genes that were functionally associated with the cell cycle or proliferation according to either KEGG or MsigDB databases (*n* = 28). This resulted in a list of 389 unique murine genes, which were mapped to their human orthologs using R package `BiomaRt`. The details of this procedure including cutoffs and group comparisons are also summarized in S2 Table.

To define the proliferation signatures, we used the list of genes that are functionally associated with proliferation as provided by Ben-Porath *et al* (*n* = 326) [48]. To infer activity of epithelial-to-mesenchymal transition (EMT) and mesenchymal-to-epithelial transition (MET), we selected TFs associated with either of the two processes, *SNAI1* and *SNAI2* for EMT, and *GRHL1-3* for MET. Then, we combined the selected TFs with their protein interactors and co-expressed genes according to STRING-DB (v.11.0) [49] resulting in a list of *n* = 51 genes. The resulting gene lists are summarized in S3 Data File.

### Calculation of signature intensity metric

To compare the degree of reactivation of the stemness program between different tumors, we introduced a metric called *signature intensity*, which represents the percentage of samples that belong to the Stem cluster among all samples of the given tumor. Recall that a sample was assigned to the Stem cluster (respectively, Normal cluster) if it was located closer to (respectively, further from) ESC/iPSCs according to PC1 than the global median of the PC1 axis. That is, the signature intensity *I*_*j*_ of the tumor *j* was computed as 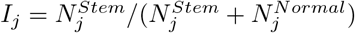 where 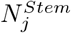 is the number of samples of the tumor type *j* in the Stem cluster and 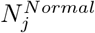 is the number of samples of the tumor type *j* in the Normal cluster.

### Survival analysis

To test whether the reactivation of the stemness program correlates with poorer prognosis, we stratified tumor samples into two clusters relative to the median PC1 value of a given tumor type. For a given tumor we subdivided samples into Stem and Normal clusters based on the proximity to iPSCs on PC1. However, instead of using the global median PC1 over all tumors as a threshold, we used the median PC1 of each tumor type. This approach resulted in tumor-specific clusters of balanced size, which were used for survival analysis. We fitted Cox-regression using Stem and Normal clusters as predictors for each tumor when clinical data were available. Hazard ratios (HR) between two clusters were then computed for all tumors using R package `survminer`.

### Statistical analysis

All statistical analyses were done using R software version 3.6.0. Confidence intervals for proportions were computed using a 2-sample z-test without continuity correction. All tests were carried out at the 5% significance level with Benjamini-Hochberg correction for multiple testing.

## Results

### Stemness genes

Several lists of stemness marker genes currently exist, however there is no universal such list [50]. For instance, a recent meta-analysis of stemness genes from several independent studies revealed that only one gene was common among lists derived from three SC populations [51]. Complementary to this, here we assembled a list of pluripotency markers using the data from a range of different experimental approaches including tissue atlases, ESC differentiation time-series, and knockdowns of pluripotency-associated transcriptional factors [41–44]. To distinguish between stemness and other traits, we removed proliferation-related genes and tissue-specific genes [47] (see Methods for details) and obtained a list of (*n* = 389) stemness genes including master regulators of stemness *POU5F1, SALL4, NANOG* as well as many other genes involved in the transcriptional regulation and cellular metabolism (S3 Data File) [52]. The comparison of this set with the ESC signatures provided in other studies [48, 53, 54] revealed a moderate intersection (Fig 1), however not as large as the intersection of ESC signatures with each other. For instance, the gene set of Bhattacharya *et al* overlapped by more than a half with the ESC signature identified by Ben-Porath *et al* (*n* = 50), while the largest intersection of our set was with the gene set of Wong *et al* (*n* = 45).

**Fig 1.**
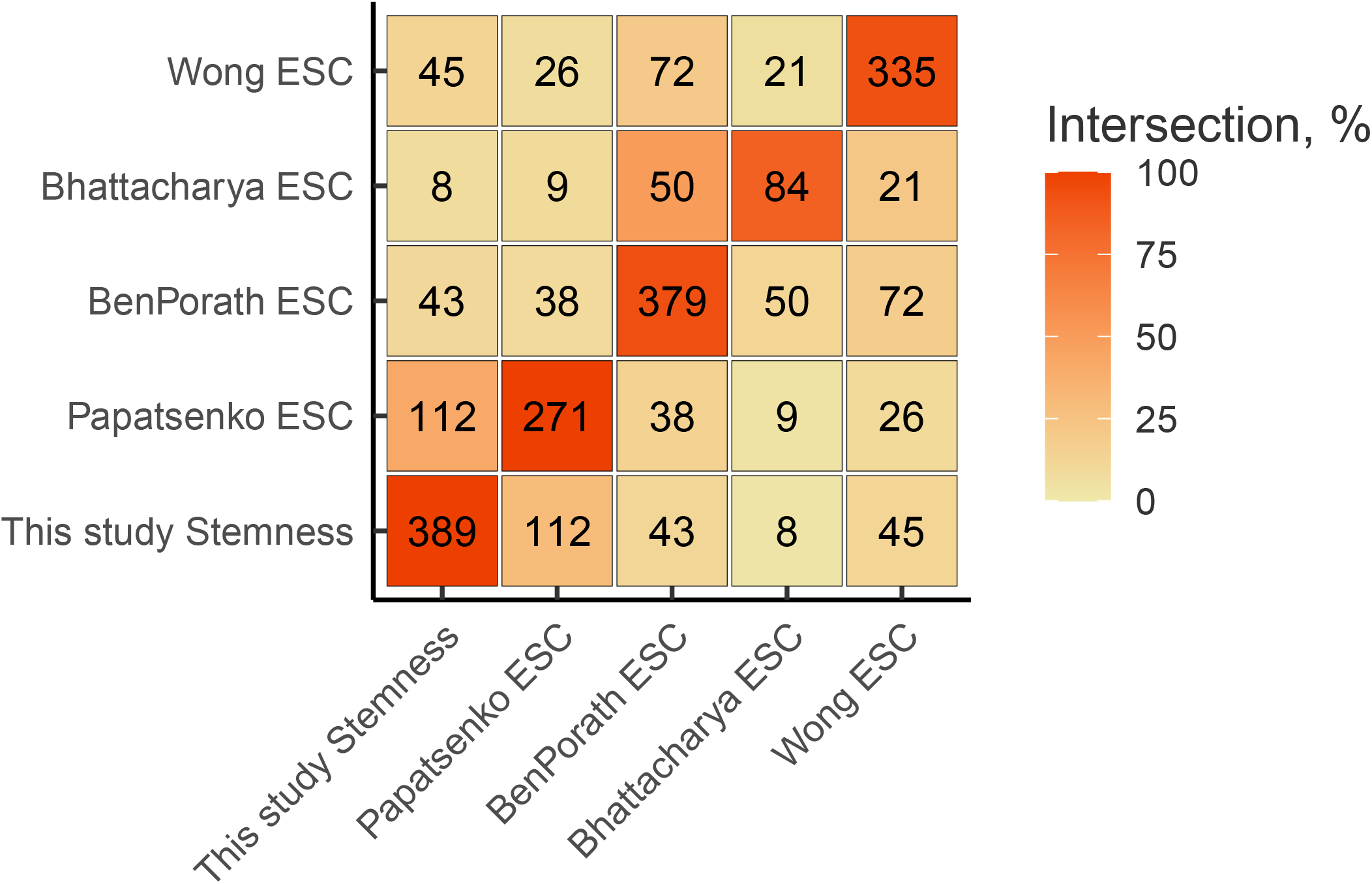
The stemness signature gene set intersects with previously published ESC signature gene sets. The number of genes in the intersection between sets on the X and Y axes is shown in the cells of the matrix. The color scale encodes the size of the intersection relative to the size of the dataset on the Y axis (above the diagonal) and relative to the size of the dataset on the X axis (below the diagonal).

Since tumors are composed of heterogeneous populations of cell types, an admixture of normal cells into tumor samples can influence global gene expression profiles and dilute stemness signature with the signal from normal cells, in which the stemness program is silent. An estimate for the cellular composition of a tumor sample is the so-called tumor purity, which can be quantitatively measured as the percentage of true tumor cells among multiple other cell types that constitute the tumor [39]. Several approaches have been developed to estimate this metric [39, 55–58]. They were recently applied to the TCGA dataset to compute the consensus tumor purity estimate (CPE), which represents the median of four different tumor purity estimates [39]. The distribution of CPE varies greatly both between and within different tumor types (S1 Fig). To account for the effect of cellular composition bias, we built a linear model (see Methods) of gene expression as a function of CPE and used the residuals of this model instead of the raw gene expression values (S2 Fig).

### Tumors span continuously from normal tissues to iPSC

To investigate how tumors, normal tissues, ESC, ASC, and iPSCs compare in terms of stemness signature, we used principal component analysis (PCA) of the adjusted gene expression profiles restricted to the set of stemness genes (Fig 2A). Tumor samples distributed over the first principal component (PC1, 24% of the total variance) forming two distinct clusters located between normal tissues and ESCs/iPSCs, while ASCs were located close to the normal tissues on the PC1 axis. At that, the iPSCs differentiation time series also clustered along PC1 so that less differentiated cells were located further from the center. Remarkably, the second principal component (PC2, 8% of the total variance) separated iPSCs and ESCs. The higher-order principal components did not show any clear separation related to stemness genes (S3 Fig), indicating that stemness signature is encoded within PC1. A similar clustering by stemness signatures from previously published gene sets did not provide a clear separation of tumors, normal samples and SCs (S4 Fig).

**Fig 2.**
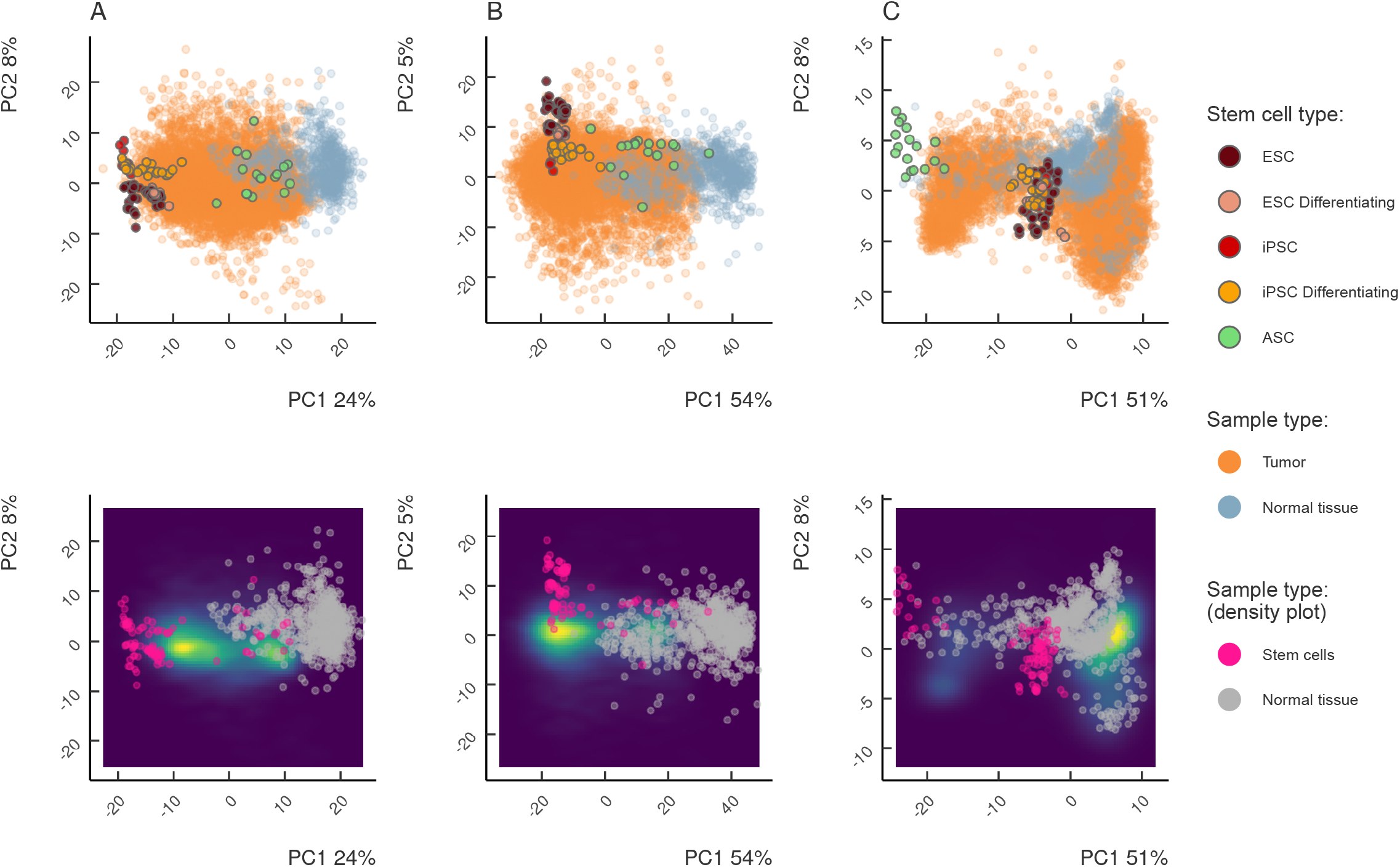
Principal Component Analysis (PCA) of the expression of stemness signatures (A), proliferation signatures (B), and genes representing EMT-MET (C) in differentiating iPSCs, ESCs, ASCs, tumors and normal tissues shown as individual samples (top) and density (bottom). Tumors span continuously between normal tissues and iPSCs in the expression subspaces of stemness, proliferation, but not EMT-MET genes. Tumor samples form two clusters: one closer to ESCs and iPSCs, another closer to the normal tissues and ASCs.

The expression of particular stemness markers followed the pattern of clustering along PC1 (S5 Fig). *SALL4*, one of the major regulators of pluripotency, was among the genes with the highest loadings in PC1, with a gradual increase in expression along the PC1 axis reaching its maximum at iPSCs. On the other hand, *SALL1*, which is suppressed in breast cancer [59], was ubiquitously downregulated in tumors, while being steadily expressed in both stem cells and normal tissues. The expression of *POU5F1*, a key regulator of stem cell pluripotency, substantially increased, while the expression of *CBX7*, a gene that encodes a Polycomb protein which globally regulates cellular lifespan [60] and is a tumor suppressor in both mice and humans [61], decreased towards ESC/iPSCs (S5 Fig).

To check whether the observed clustering along PC1 axis was consistent with patterns reported for particular tumor types, we analyzed the stemness signature in two related, yet different cancers, lung squamous cell carcinoma (LUSC) [62] and lung adenocarcinoma (LUAD) [63]. LUSC has a higher mortality rate compared to LUAD and shows a pronounced upregulation of *sonic-hedgehog*, a major regulator of the developmental pathway linked to stemness and proliferation, which is often active in adult stem cells and is absent in LUAD [64, 65]. Consistent with this, LUSC was located closer to SC along the PC1 axis, while LUAD was closer to normal tissues (S6A Fig). In line with previous observations, we observed an increased expression of the pluripotency-associated transcription factor SOX2 and reduced expression of CBX7 in LUSC (S6B,C Fig). Additionally, genes differentially expressed in LUSC were enriched for development-associated GO terms (S7 Fig).

### Analysis of proliferation and EMT signatures

The expression of stemness genes revealed clustering of tumor samples in between normal tissues and ESC/iPSCs, which may not be a unique property of stemness genes. However, no clustering was observed when we performed PCA on a random set of genes that were matched by expression levels to stemness genes (S8 Fig). In contrast, when we repeated the same analysis using proliferation markers, tumor samples again scattered across the PC1 axis (55% of variance) forming two separate clusters (Fig 2B), and the differentiation states of iPSCs separated concordantly along this axis. As before, PC2 separated ESCs from iPSCs, while no clear separation was observed in higher order principal components suggesting that PC1 represents the proliferation signature (S3B Fig).

In contrast, the clustering of tumors, normal tissues, and iPSCs with respect to the epithelial to mesenchymal transition (EMT) signature was drastically different (Fig 2C). Except for the marginal standing of ASC, in which EMT must be active, we did not observe any clear separation of tumor samples from normal tissues, nor did we detect a directional trend of tumor samples along any axes. There was no clear separation in higher-order principal components, and most of the tumors showed epithelial phenotype similar to their tissues of origin, except for hepatocellular carcinoma, melanoma and brain tumors (S3C Fig). Instead of the gradient of EMT signature, we observed a switch-like state of master regulators of EMT (S9 Fig). Altogether, this analysis failed to capture EMT signature in tumor tissue, possibly because cells undergoing EMT are located at metastatic outgrowth or even dispersed as a transient circulating population thus preventing their detection in bulk sequencing [66].

### Heterogeneity of stemness and proliferation signatures across tumor types

The analysis of stemness and proliferation signatures revealed two clusters of tumor samples, one of which was located closer to iPSCs along PC1, and the other was located closer to normal tissues (Fig 2A-B). Since this separation occurred along the PC1 axis in both signatures, we formally defined the clusters by their position relative to the median of the PC1 across all tumor samples (Fig 3, black vertical line). Namely, samples located to the right of the median of PC1 axis towards normal tissues were assigned to the *Normal* cluster, while samples located to the left of the median towards iPSC were assigned to *Stem* cluster.

**Fig 3.**
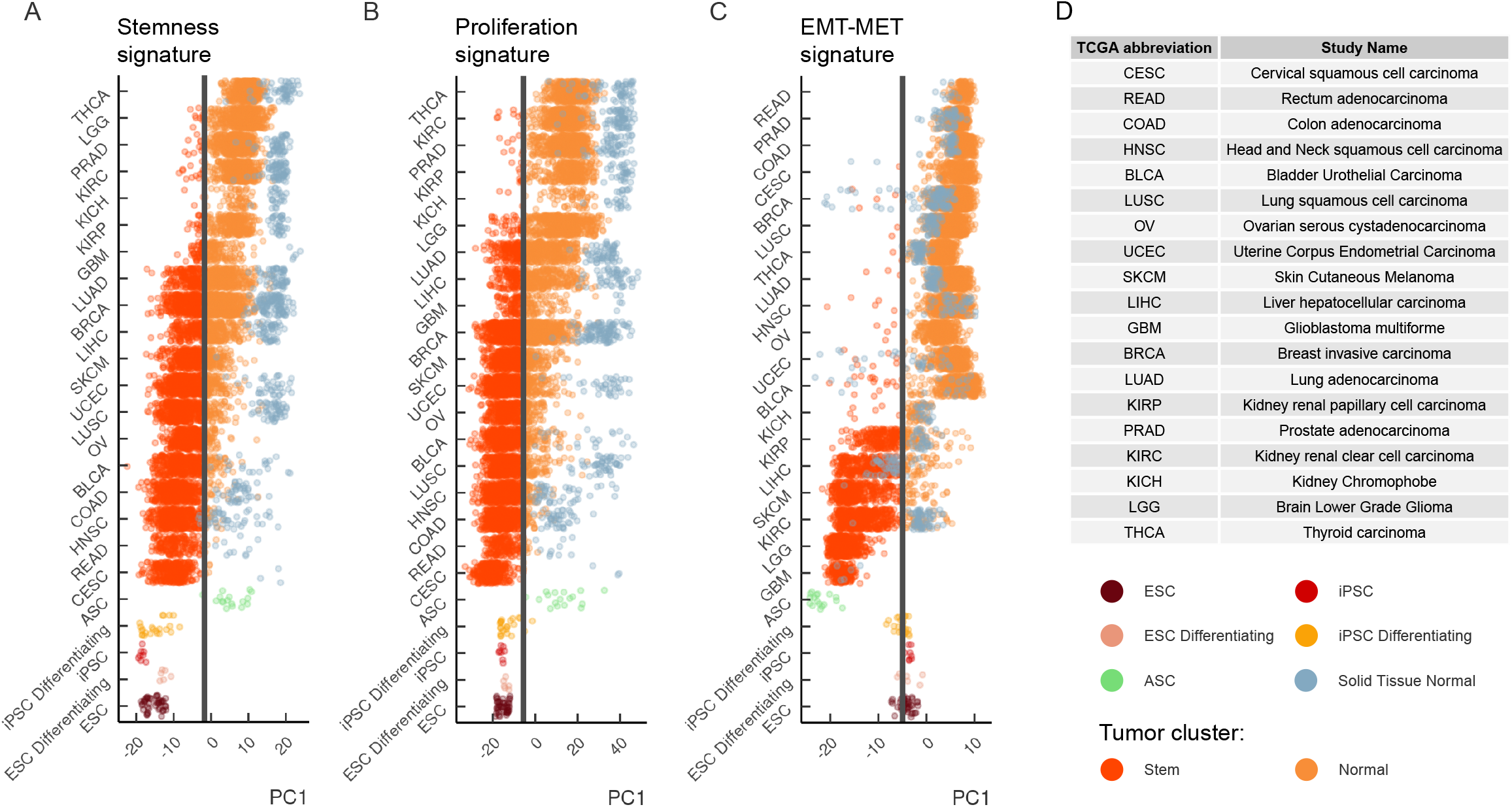
Heterogeneity of stemness and proliferation signatures across tumor types. The first principal component (PC1) encodes the signature of stemness (A), proliferation (B), and EMT-MET signature genes (C). The samples are plotted separately for each tumor in the TCGA project, normal tissues, and iPSCs. Tumor samples are colored according to their position relative to the PC1 median (black vertical line). Tumor samples located to the right (left) of the median are assigned to the Normal (Stem) cluster, respectively. (D) The standard TCGA notation for cancer types.

Next, we investigated whether any tumor type was specifically enriched in one of the two clusters (Fig 3A-C). Indeed, some tumors were fully localized to one of the clusters, e.g. Rectal Adenocarcinoma and all subtypes of Renal Cell Carcinoma (KIRP, KIRC, KICH), while others showed a significant heterogeneity in terms of both stemness and proliferation signatures, e.g. Lung Adenocarcinoma and Liver Hepatocellular Carcinoma. To quantify the heterogeneity of stemness and proliferation signatures within each tumor type, we computed a metric called *intensity*, which is defined as the proportion of samples of a given tumor type that are located in the Stem cluster (see Methods for details). In other words, this metric captures the degree, to which each tumor type collectively expresses stemness or proliferation signatures.

Despite stemness and proliferation intensities being strongly correlated, they differ significantly for some tumor types (Fig 4A, B). For instance, the stemness intensity of glioblastoma (GBM) is 28% *±* 7%, while its proliferation intensity is 50% *±* 8%. Conversely, the stemness intensity of hepatocellular carcinoma (LIHC) is 46% *±* 5%, while its proliferation intensity is 31%*±* 5%. Thus, these two metrics capture consistent, yet different aspects of tumor gene expression. Most tumors with high purity have low stemness and proliferation intensities, consistent with lower aggressiveness of such tumors [39], while tumors of higher stage tend to have higher intensity of both signatures.

**Fig 4.**
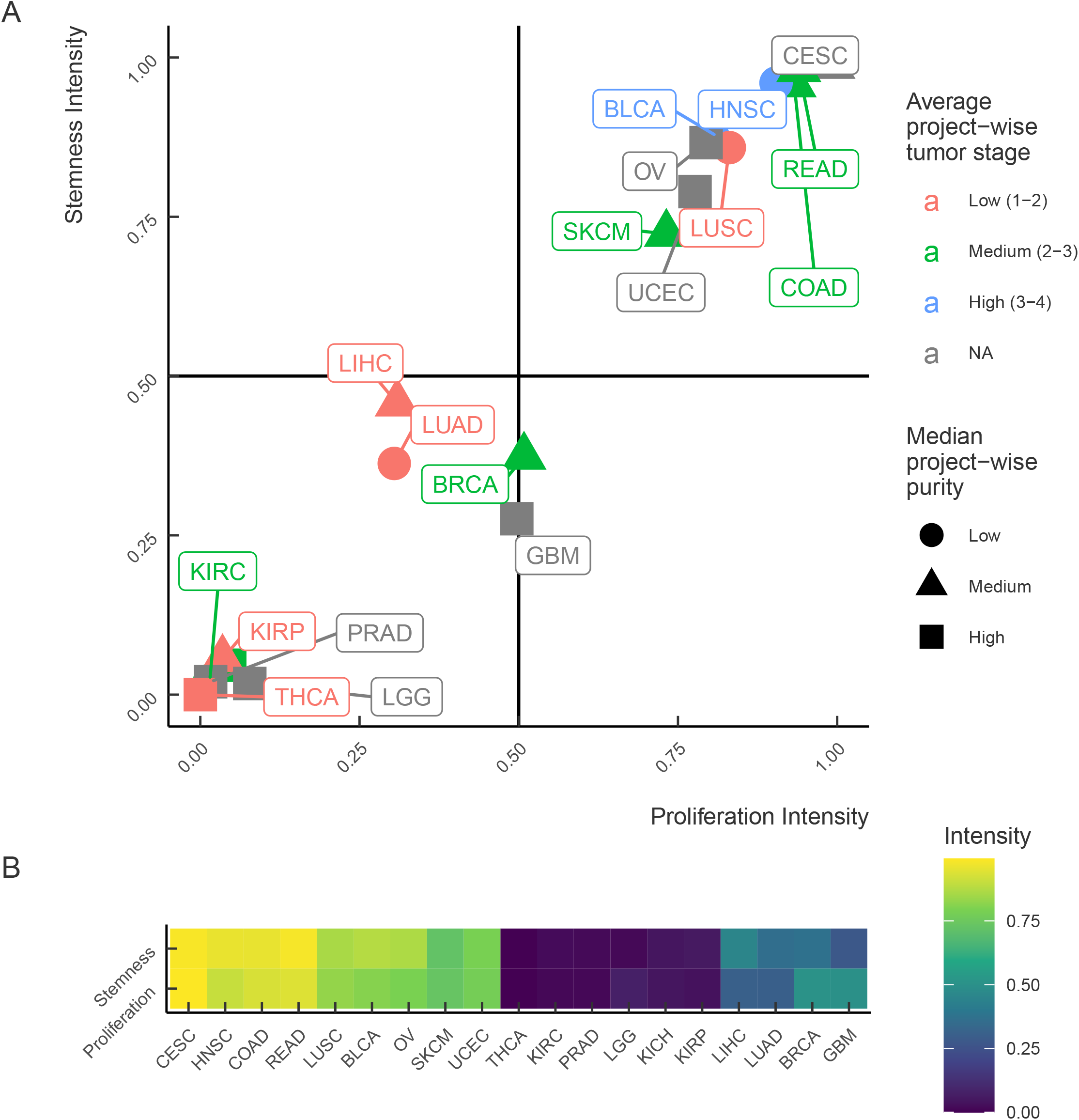
The relationship between stemness and proliferation intensities. (A) Stemness intensity vs. proliferation intensity in 19 solid tumors with different average project-wise tumor stage and median project-wise purity. (B) A heatmap of stemness and proliferation intensities across 19 solid tumors from TCGA. Two groups of tumor samples are identified: top-right quadrant (stemness Intensity*>* 0.5 and proliferation Intensity*>* 0.5) and bottom-left quadrant (stemness Intensity*<* 0.5 and proliferation Intensity*<* 0.5). The notation for cancer types as in Fig 3.

### Intertumor variability correlates with survival

The intensity of stemness and proliferation signatures separates most tumors into two groups corresponding to Normal and Stem clusters by the median of the PC1 axis. However, this subdivision reflects global heterogeneity of stemness among all analyzed tumor types. In order to assess the intertumor heterogeneity, i.e., the heterogeneity between different tumors of the same tumor type, we used the PC1 median of a given tumor type rather than the global PC1 median across all tumor samples to cluster samples into Stem and Normal groups specific to each tumor type (Fig 5A). This approach provided balanced groups, which we used for survival analysis.

**Fig 5.**
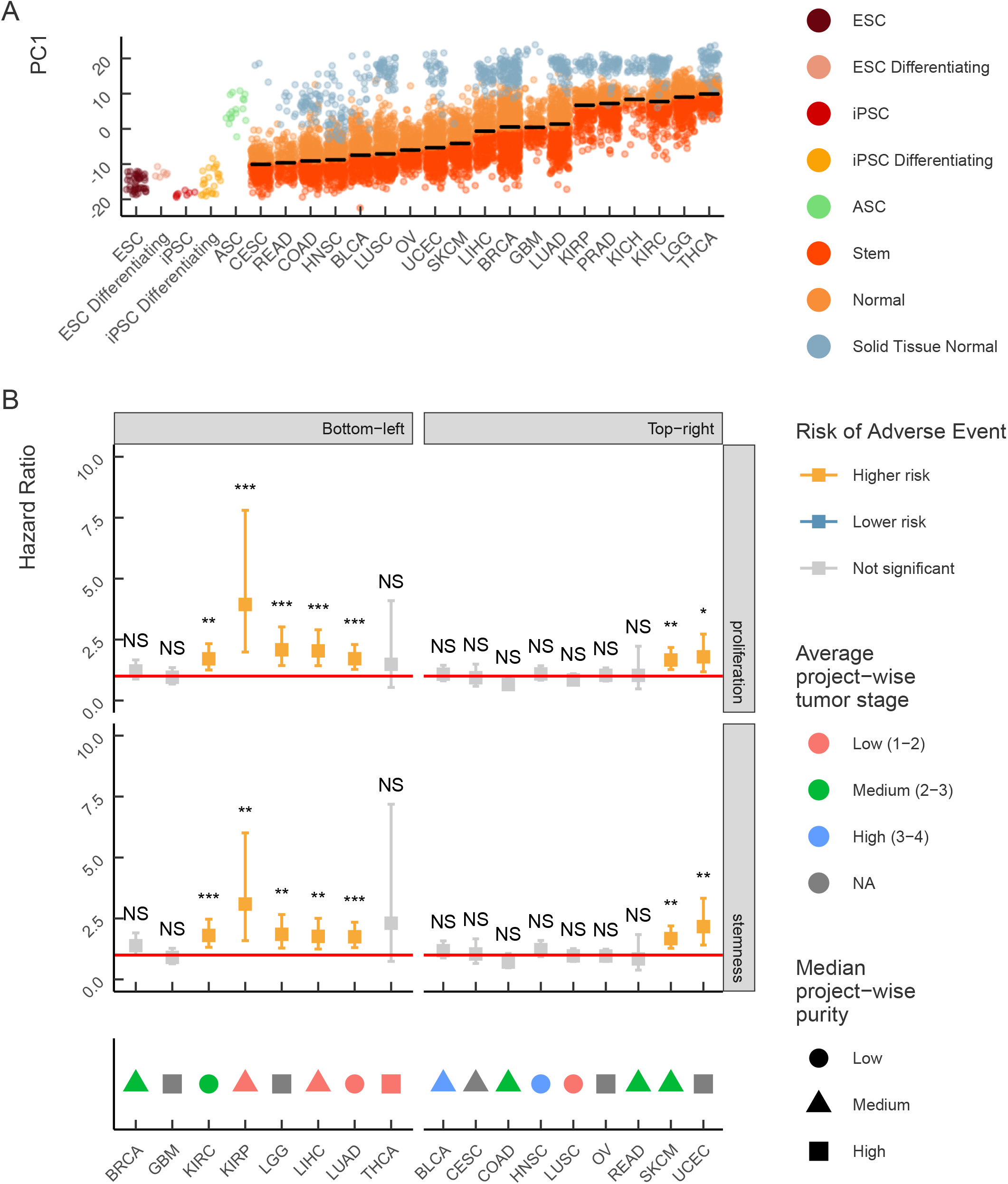
Intertumor heterogeneity of stemness and proliferation signatures. (A) Stem and Normal clusters specific to each tumor type are defined by the location of a sample relative to the median PC1 value across all samples of the given tumor (plotted as horizontal black lines). Samples above (below) the median are assigned to the Normal (Stem) cluster, respectively. (B) Hazard ratios for the comparison of patient survival in Stem and Normal clusters in two sample groups (top-right and bottom-left in Fig 4) with respect to stemness and proliferation intensity. Error bars denote 95% confidence intervals. Tumor types are annotated at the bottom according to the average tumor stage and the median tumor purity. Codes for statistical significance reported are as follows: NS − p-value *>* 0.05, ∗ − 0.01 *<* p-value *<* 0.05, ∗∗ − 0.001 *<* p-value *<* 0.01, ∗ ∗ ∗ − p-value *<* 0.001. The notation for cancer types as in Fig 3.

We observed a significant difference in survival between the Stem and Normal clusters only in a fraction of tumors (Fig 5B). Remarkably, the hazard ratio was significant for tumors that did not show the highest stemness and proliferation intensity (the bottom left quadrant in Fig 4), while tumors with the highest stemness and proliferation intensity (the upper right quadrant in Fig 4) did not show a significant difference in survival. For instance, we did not observe a significant difference in hazard ratio for LUSC despite high stemness and proliferation intensity, while skin cutaneous melanoma (SKCM), which had the stemness intensity on the level of LUSC, showed a significant difference in the survival between Stem and Normal clusters (Fig 6). A similar heterogeneity was observed for tumors with low stemness intensity such as kidney renal carcinoma (KIRC) and low grade glioma (LGG).

**Fig 6.**
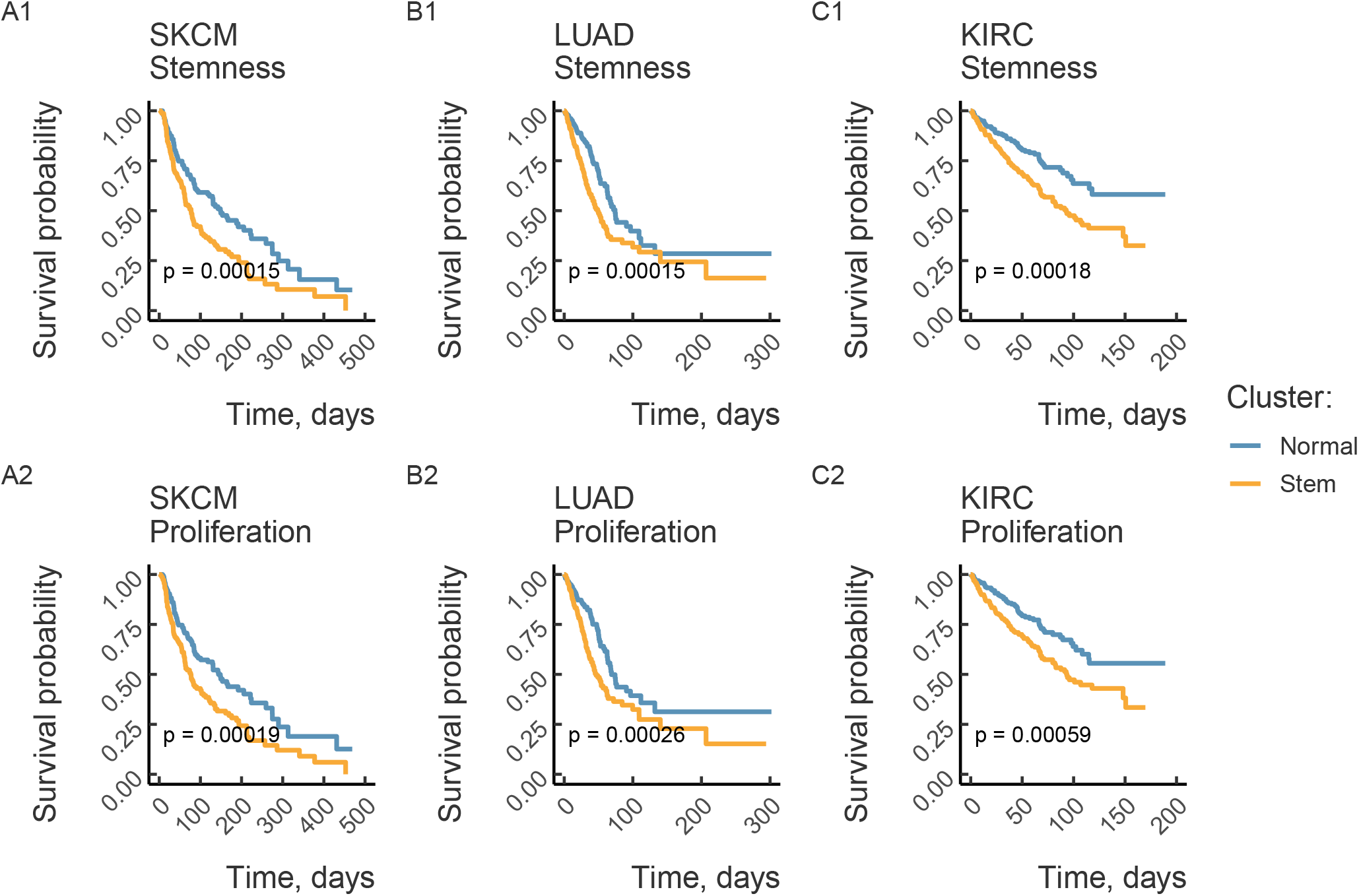
Intertumor heterogeneity of stemness and proliferation is predictive of poor survival. Kaplan-Meier survival curves grouped by clustering according to stemness (proliferation) signature are plotted for skin melanomas (A), lung adenocarcinoma (B) and renal clear cell carcinoma (C). P-values from the logrank test are reported. The notation for cancer types as in Fig 3.

### Intratumor heterogeneity

Unlike intertumor variability, which reflects differences between different tumors of the same type from different patients, intratumor variability refers to genotypic and phenotypic differences between clonal populations of cells within a tumor. The quantitative measurement of the transcriptional diversity of cells within a tumor are provided by transcriptomic profiling at a single-cell resolution [67]. To assess the intratumor heterogeneity of stemness signature, we re-analyzed the transcriptomes of 65,362 unsorted single cells from metastatic colorectal cancers [32]. We used the loadings, i.e., the coefficients of the linear combination of stemness signature genes, that correspond to the PC1 axis in Fig 2A to project the transcriptional profiles of single cancer and non-cancer cells onto PC1 (Methods). The 2D representation using uniform manifold approximation and projection (UMAP) [68] revealed a clear separation of cancer and non-cancer cells consistent with the annotation from [32] (Fig 7). Remarkably, the cancer cell cluster shows a considerable increase of the stemness signature encoded within PC1 (Fig 7A). However, not only the absolute value of the stemness signature, but also its variability was higher in the cancer cell cluster compared to non-cancer cell cluster indicating that the heterogeneity of stemness signature is observed at all levels of cancer organization, including single-cell level.

**Fig 7.**
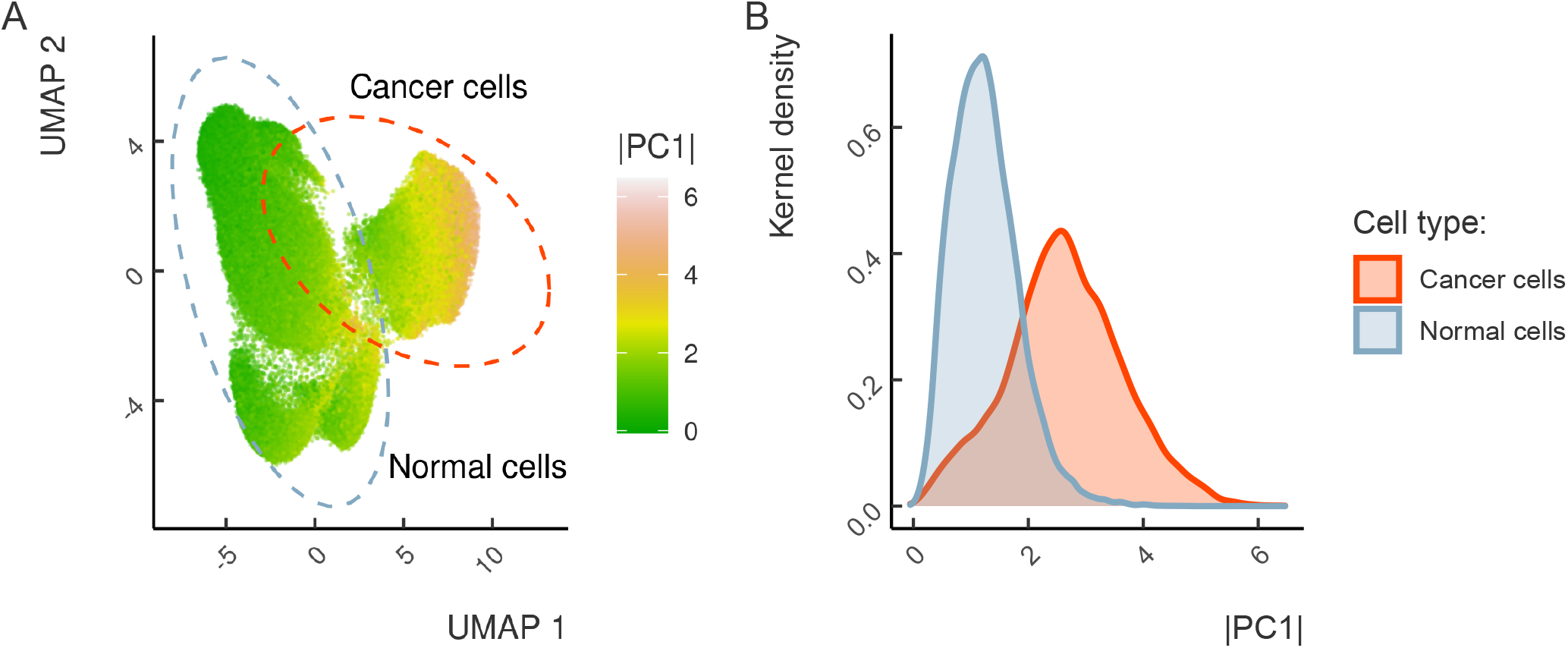
Cancer cells show substantial intratumor heterogeneity of stemness. (A) UMAP dimensionality reduction of the colorectal cancer single-cell RNA-seq dataset. The annotation of normal and cancer cell clusters is from [32]. The cells are colored by the degree of stemness signature reactivation (PC1). (B) The distribution of stemness signature (PC1) (X axis) in normal and cancer cells.

## Discussion

Transcriptome comparisons have been a powerful tool for studying gene expression signatures in disease [69–71]. We used principal component analysis (PCA), which represents each sample as a point in an *n*-dimensional space with coordinates corresponding to gene expression values and applies a dimensionality reduction to identify principal components with the largest variance. The outcome of these transformations, however, critically depends on the set of genes that were chosen initially. For instance, the use of different gene sets led to opposite outcomes with tissue-dominated and species-dominated clustering [72]. In this work, we identified a gene set corresponding to the stemness signature encoded in PC1, which followed the stemness gene expression gradient from the normal tissues through tumors and ASC to ESC and iPSCs. This gene set provides a better representation of the stemness axis compared to previously published gene sets [48, 53, 54].

The TCGA dataset showed a consistent reactivation of stemness in all solid tumors in comparison to normal tissues, in accordance with other reports [73–75]. At the same time, the degree of reactivation of the stemness and proliferation signatures varied greatly between different tumor types and also inter- and intratumorally, thus extending the results of previous studies on stemness heterogeneity to the TCGA dataset [27, 76]. Remarkably, tumors originating from the tissues that actively interact with the outside environment (lungs, urinary system, skin, intestines) show strong reactivation of both stemness and proliferation programs consistently with the hypothesis that tumors inherit their self-renewal capacity from the tissues of origin.

An important aspect of the reactivation of gene expression programs in tumors is formulated by the Lineage Addiction Model, which suggests that the mechanisms that promote tumor progression involve master regulatory genes that also exert key survival roles in development [77]. Multiple examples of cancer lineage addiction have been reported [78–81], however the effect of lineage addiction on the reactivation of the stemness program remains to be studied in detail. In our analysis, only a few tumor types (Breast Cancer, Lung Adenocarcinoma, Lung Squamous Cell Carcinoma) spanned the entire PC1 axis, while most tumor types were fully localized to either Stem or Normal cluster. This observation supports the idea that the majority of tumors are restricted to specific phenotypic spaces with respect to the reactivation of the stemness program.

In the study of Ben-Porath *et al*, high expression of stemness genes was shown to be predictive of poor survival among the patients of three breast cancer cohorts [48]. Other studies also reported reduced survival for the tumors of stem phenotype [82–85]. Overall, as one of the hallmarks of cancer [86], stemness has been suggested before as a universal predictor of patient survival [27]. In our analysis, however, only seven out of sixteen tumors, for which patient survival data were available, showed a significant negative correlation of stemness with survival. These results highlight the absence of universal ties between patient survival and the degree of stemness signature reactivation, indicating a substantial degree of heterogeneity in tumor reaction to the latter [26]. This analysis suggests that in spite of associations with tumor stage and aggressiveness, the stemness is not a universal predictor of survival and could be regarded instead as a function of the molecular profile that is specific for every tumor type.

The major limitation of the present study arises from the use of bulk RNA-seq, which is unable to capture cellular heterogeneity. Single-cell RNA-seq experiments provide an orthogonal view to the bulk RNA-seq by measuring the transcriptional profiles of individual cells, however at the expense of sparse coverage. Here, we used colorectal cancer to demonstrate as a proof of principle that the expression of stemness signature is highly heterogeneous not only intertumorally, but also at the level of individual cancer cells. Additionally, it was reported that specific patterns of hyper/hypo methylation follow the stem phenotype in cancer cells [87, 88]. Therefore, another source of stemness heterogeneity may come from the epigenetic component or from other factors such as variability in pre-mRNA splicing [89] and expression of non-coding genes, e.g. transposons [90]. A combined analysis of single-cell transcriptomes, pre-mRNA splicing by bulk RNA-seq, epigenetic assays based on ChIP-seq, and non-coding RNA quantification is needed for the detailed characterization of the stemness heterogeneity landscape. Growing amounts of this information in the public domain enable such characterization from the multi-omics perspective and open new directions for future studies.

## Conclusions

Tissue-independent reactivation of the stemness program represents a unifying feature which ties together tumors of different origins. Nonetheless, the degree to which this program becomes reactivated strongly fluctuates between different tumor types and also inter- and intratumorally, thus suggesting that the effect of the stemness program on the tumor phenotype is highly heterogeneous. Multiple studies, including those based on single cell technology, have described tumor heterogeneity arising at different levels through various mechanisms. Here, we pinpoint yet another and previously unappreciated aspect of heterogeneity, one that is related to the reactivation of the stemness program, thus adding to an already complex picture of tumorigenesis and potentially impacting the diagnostic protocols and the development of new anticancer treatments.

## Supporting information

**S1 Fig. Consensus purity estimate (CPE)**. The distribution of the consensus purity estimate (CPE) values across the 19 tumors. The tumor types on the *x* axis are listed by in the descending order by the mean CPE value.

**S2 Fig. Correction of gene expression values**. (A) The distribution of log_2_(1 + *CPM*), where CPM denotes median counts per million, for the raw and

CPE-corrected gene expression values. The gene expression counts were first normalized by edgeR and then corrected for tumor purity (See Methods - Correction for tumor purity). (B) A scatter plot of CPE-corrected vs. raw log_2_(1 + *CPM*). Each point represents a gene.

**S3 Fig. Clustering by higher order principal components**. Higher order principal components corresponding to Fig 2. Panels (A), (B), and (C) correspond to the stemness, proliferation, and EMT-MET signatures, respectively.

**S4 Fig. Clustering according to previously reported stemness signatures**. Clustering of tumor, normal, and ESC samples according to previously reported stemness signatures: (A) – Ben-Porath *et al* [48], (B) – Bhattacharya *et al* [53], (C) – Wong *et al* [54].

**S5 Fig. Expression of transcriptional repressors and positive regulators of stemness**. Expression of transcriptional repressors (*CBX7, SALL1*) and positive regulators of stemness (*POU5F1, SALL4*) in tumor samples along the PC1 axis in Fig 2A. Samples were stratified into quartiles according to PC1.

**S6 Fig. LUSC and LUAD tumors**. The positions of LUSC and LUAD tumors in the clustering diagram in Fig 2A. All tumor samples except for LUSC and LUAD are colored gray. The expression of CBX7 drops (B), and the expression of SOX2 increases (C) in tumors on the way from normal samples to iPSCs and ESCs.

**S7 Fig. Differentially expressed genes in LUSC and LUAD**. The differentially expressed genes in the LUSC compared to the LUAD and the respective enrichment of their associated GO-terms.

**S8 Fig. PCA clustering with respect to the control sets of random genes**. PCA clustering of tumor, normal, and ESC samples based on the controls sets of random genes that were matched by expression levels to stemness (A), proliferation (B), and EMT (C) signatures.

**S9 Fig. The gradient of expression of mesenchymal and epithelial genes with respect to EMT signatures**. The gradient of expression of mesenchymal (A-D) and epithelial (E-F) genes on the PCA clustering diagram corresponding to EMT signatures (Fig 2C). Normal tissues are shown as gray background. Stem cells are colored as in Fig 2.

**S1 Table. The RNA-seq accession numbers**. The accession numbers and description of the public RNA-seq datasets that were used in the principal component analysis along with TCGA data; *n* denotes the (effective) dataset size.

**S2 Table. The GEO accession numbers**. GEO accession numbers of the datasets that were used to assemble a set of stemness signature genes. Additional information is reported in Methods (see Signature gene sets section for details). log_2_ *FC* denotes log-fold-change threshold (Treatment vs Control); log_2_ *E* denotes log gene expression level threshold (Treatment + Control); Comparisons denotes the list of comparisons for differential gene expression analysis.

**S1 Data File. TCGA sample subtype annotation and Stem cluster attribution**

**S2 Data File. Table of literature sources of manually curated stemness markers**

**S3 Data File. The list of stemness, proliferation and EMT/MET signature genes**

## Acknowledgements

We thank Ivan Kulakovskii and Dmitrii Scherbinin for the help with curating and processing the data of Dmitrii Papatsenko. All living authors commemorate the contribution of Dmitrii Papatsenko, a cancer biologist who tragically died of a rapidly progressing lung cancer.

## Abbreviations

CSC: Cancer Stem Cells
SC: Stem Cells
ESC: Embryonic Stem Cell
ASC: Adult Stem Cell
iPSC: Induced Pluripotency Stem Cells
TCGA: The Cancer Genome Atlas
CPE: Consensus Tumor Purity Estimate
EMT: Epithelial-to-Mesenhymal Transition
MET: Mesenchymal-to-Epithelial Transition
PC: Principal Component

## Availability of data and materials

The datasets analyzed during the current study are available in the GDC Data Portal repository, https://portal.gdc.cancer.gov/repository. The datasets generated during the current study are available online at https://doi.org/10.5281/zenodo.5101514. The source code used for the analysis is available at https://github.com/melonheader/bulk-cancer-stemness.

## Authors’ contributions

DPa and DPe designed the study; DPe supervised the study; AB performed the data analysis; TI performed single-cell data analysis; DPe and AB wrote the first draft of the manuscript. DPe, AB and MG edited the final version of the manuscript. All authors have read and approved the manuscript.

## Notes

https://doi.org/10.5281/zenodo.5101514

### Competing Interest Statement

The authors have declared no competing interest.

