## Supplementary information for "Transcriptome analysis reveals high tumor heterogeneity with respect to re-activation of stemness and proliferation programs"

[1]Center of Life Sciences, Skolkovo Institute of Science and Technology, 143026 Moscow, Russia;
[2]Berlin Institute for Medical Systems Biology, Max-Delbrück-Center for Molecular Medicine in the Helmholtz Association, Hannoversche Str. 28, 10115 Berlin, Germany;
[3]Jenguro/Ingal, Ermolaevskiy lane 25, 123001 Moscow, Russia;
[†] Deceased

### Supplementary Figures


| | | |
|---|---|---|
| S1 | Consensus purity estimate (CPE) | 3 |
| S2 | Correction of gene expression values | 4 |
| S3 | Clustering by higher order principal components | 5 |
| S4 | Clustering according to previously reported stemness signatures | 6 |
| S5 | Expression of transcriptional repressors and positive regulators of stemness | 7 |
| S6 | The positions of LUSC and LUAD tumors | 8 |
| S7 | Differentially expressed genes in LUSC and LUAD | 9 |
| S8 | PCA clustering with respect to the control sets of random genes | 10 |
| S9 | The gradient of expression of mesenchymal and epithelial genes with respect to EMT signatures | 11 |


### Supplementary Tables


| | | |
|---|---|---|
| S1 | The RNA-seq accession numbers | 12 |
| S2 | The GEO accession numbers | 13 |

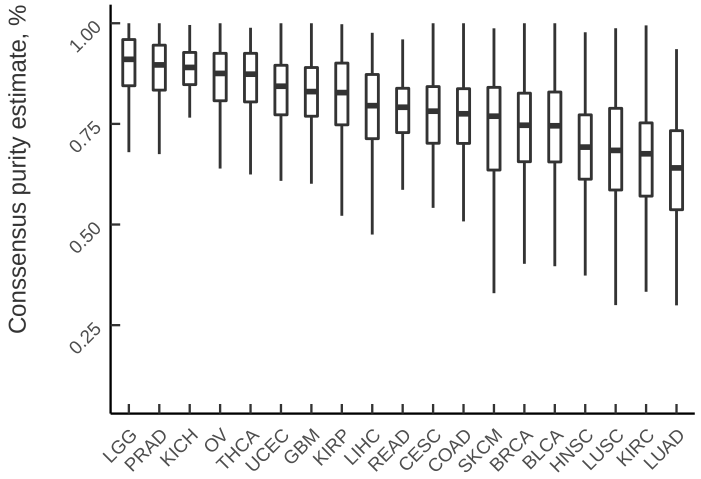


Figure S1: The distribution of the consensus purity estimate (CPE) values across the 19 tumors. The tumor types on the x axis are listed by in the descending order by the mean CPE value.

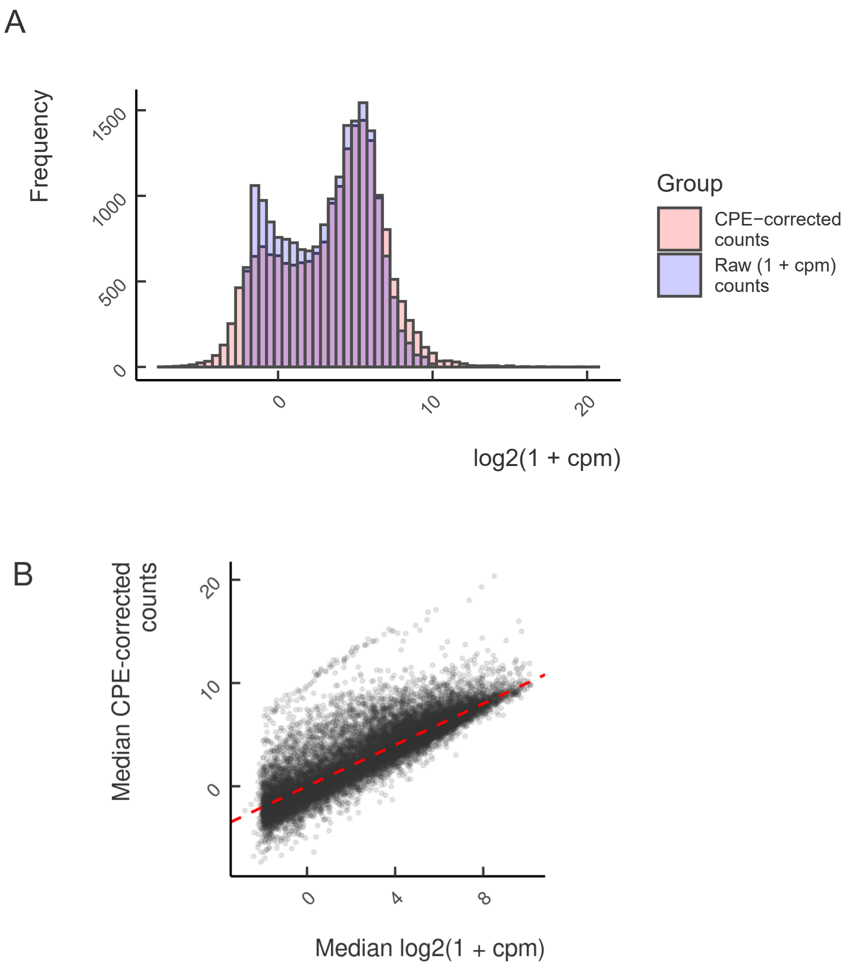


Figure S2: (A) The distribution of $\log_2(1+CPM)$, where CPM denotes median counts per million, for the raw and CPE-corrected gene expression values. The gene expression counts were first normalized by edgeR and then corrected for tumor purity (See Methods - Correction for tumor purity). (B) A scatter plot of CPE-corrected vs. raw $\log_2(1+CPM)$. Each point represents a gene.

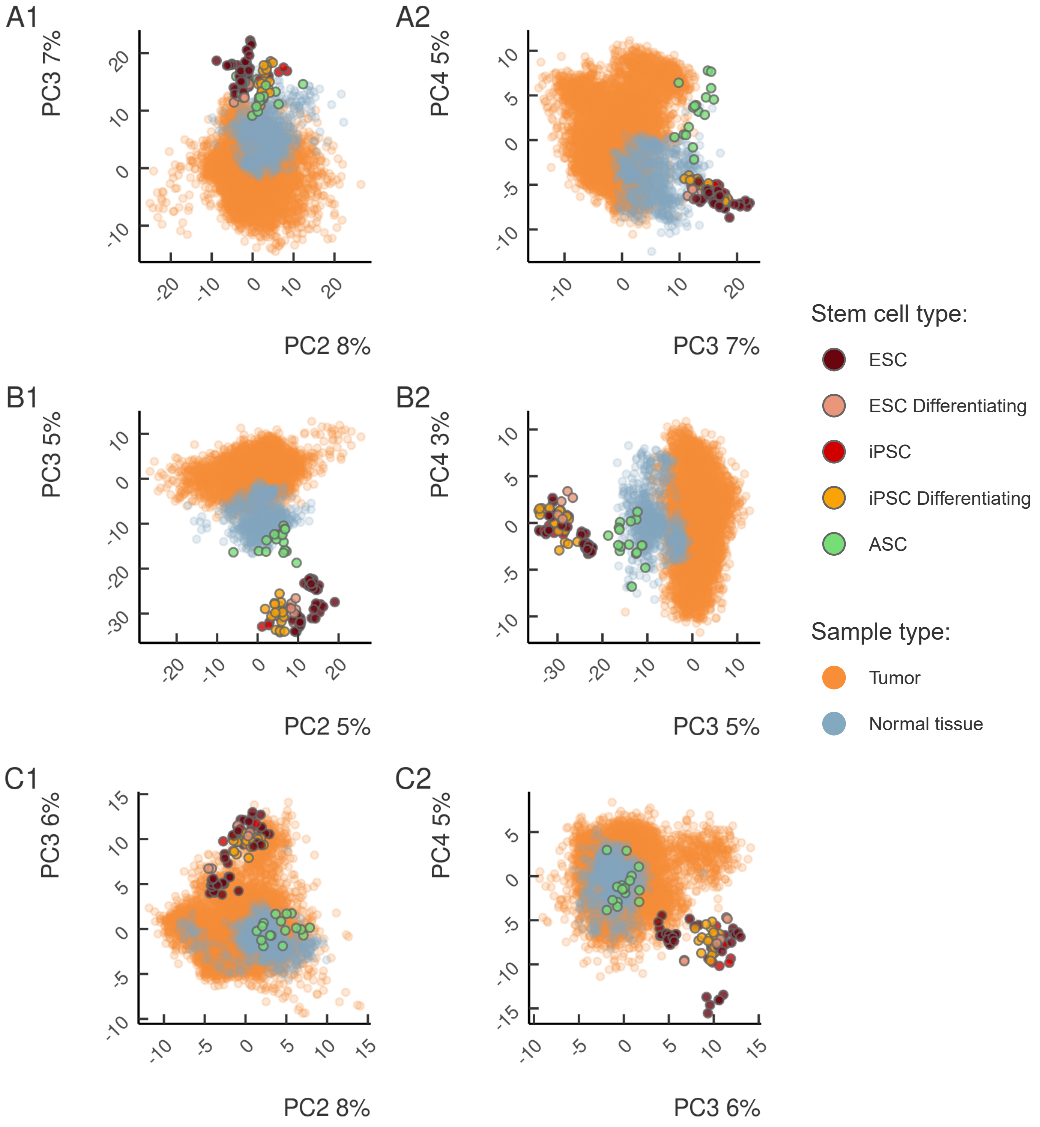


Figure S3: Higher order principal components corresponding to Fig 2. Panels (A), (B), and (C) correspond to the stemness, proliferation, and EMT-MET signatures, respectively.

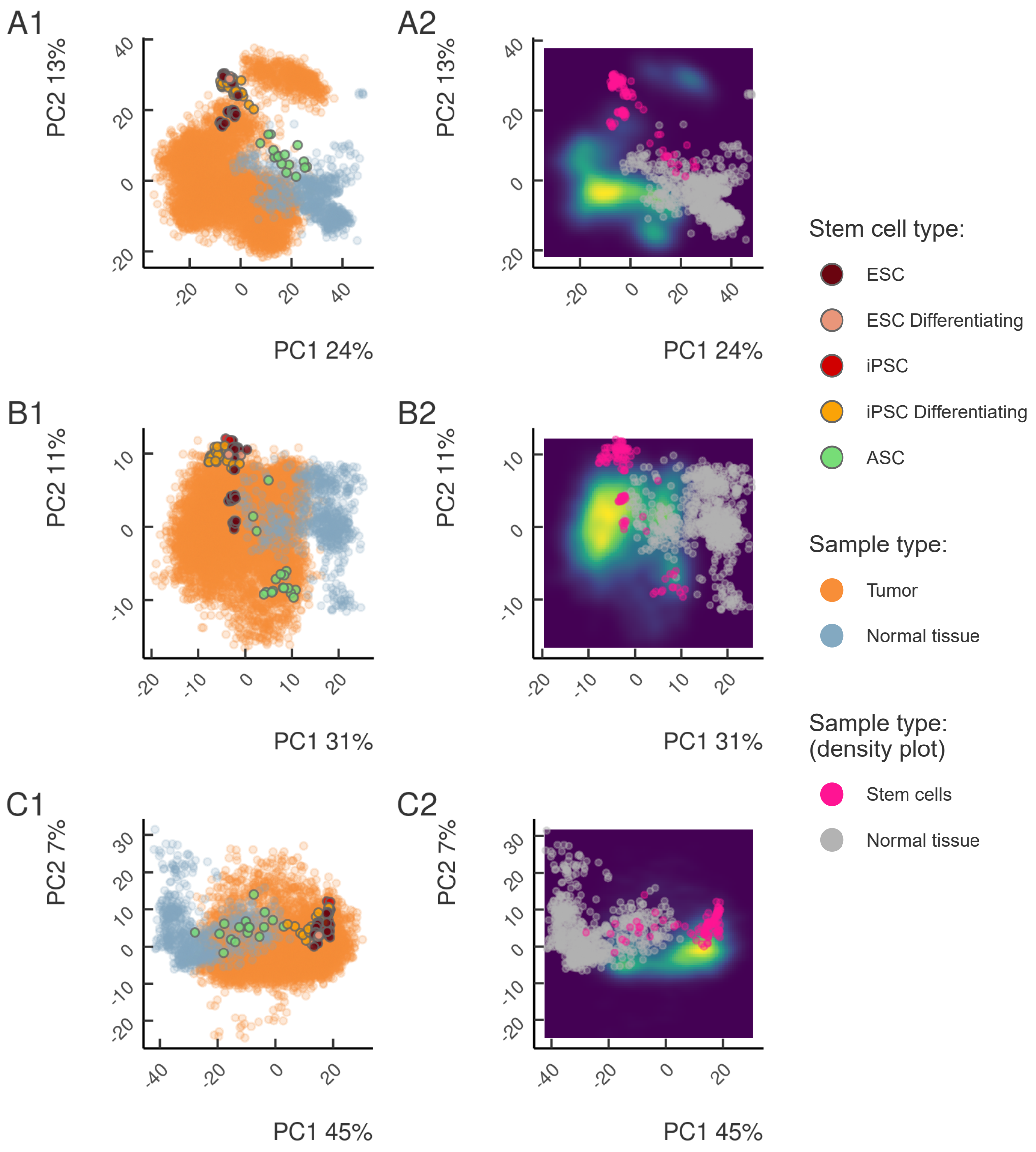


Figure S4: Clustering of tumor, normal, and ESC samples according to previously reported stemness signatures: (A) – Ben-Porath *et al* [1], (B) – Bhattacharya *et al* [2], (C) – Wong *et al* [3].

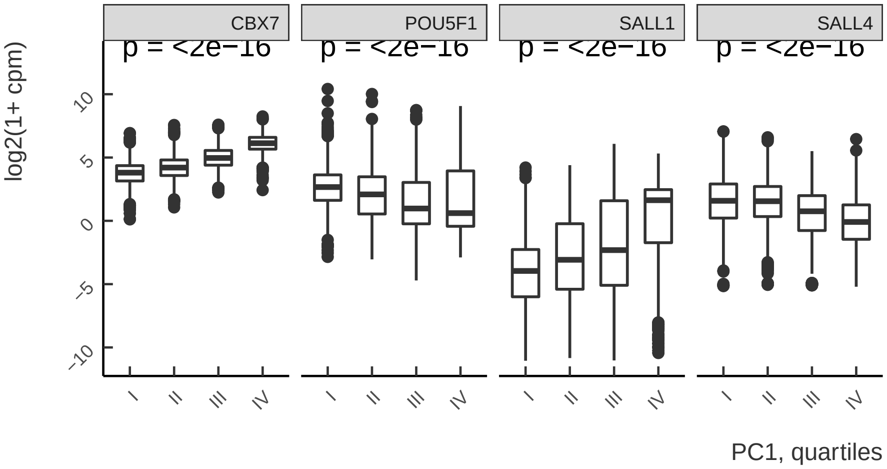


Figure S5: Expression of transcriptional repressors (*CBX7, SALL1*) and positive regulators of stemness (*POU5F1, SALL4*) in tumor samples along the PC1 axis in Fig 2A. Samples were stratified into quartiles according to PC1.

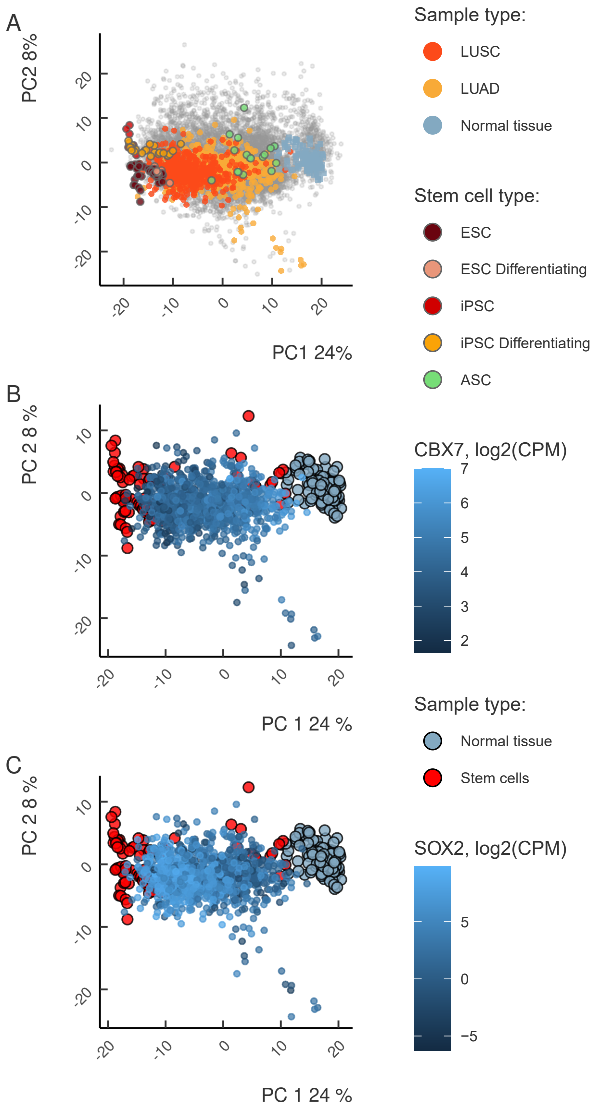


Figure S6: The positions of LUSC and LUAD tumors in the clustering diagram in Fig 2A. All tumor samples except for LUSC and LUAD are colored gray. The expression of CBX7 drops (B), and the expression of SOX2 increases (C) in tumors on the way from normal samples to iPSCs and ESCs.

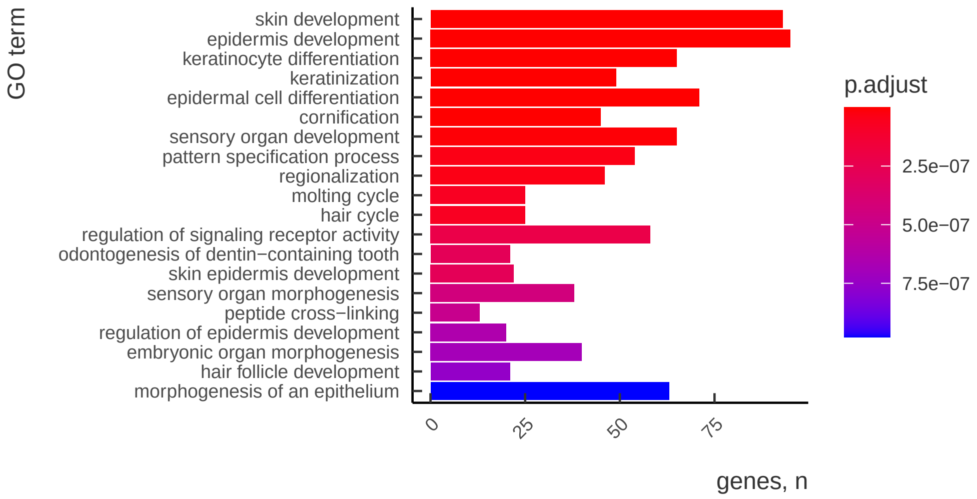


Figure S7: Differentially expressed genes (DEGs) in the LUSC compared to the LUAD and the respective enrichment of their associated GO-terms.

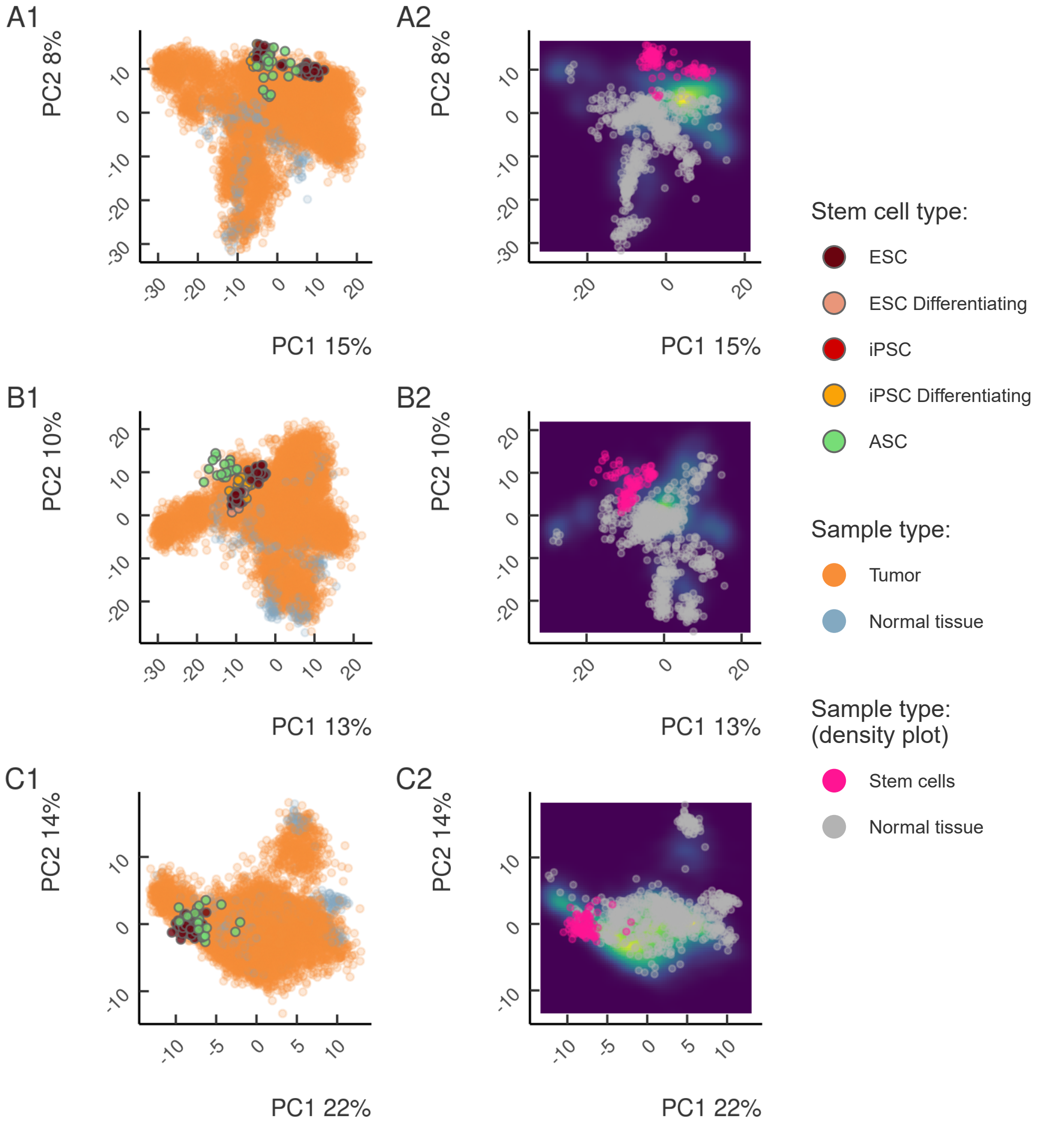


Figure S8: PCA clustering of tumor, normal, and ESC samples based on the controls sets of random genes that were matched by expression levels to stemness (A), proliferation (B), and EMT (C) signatures.

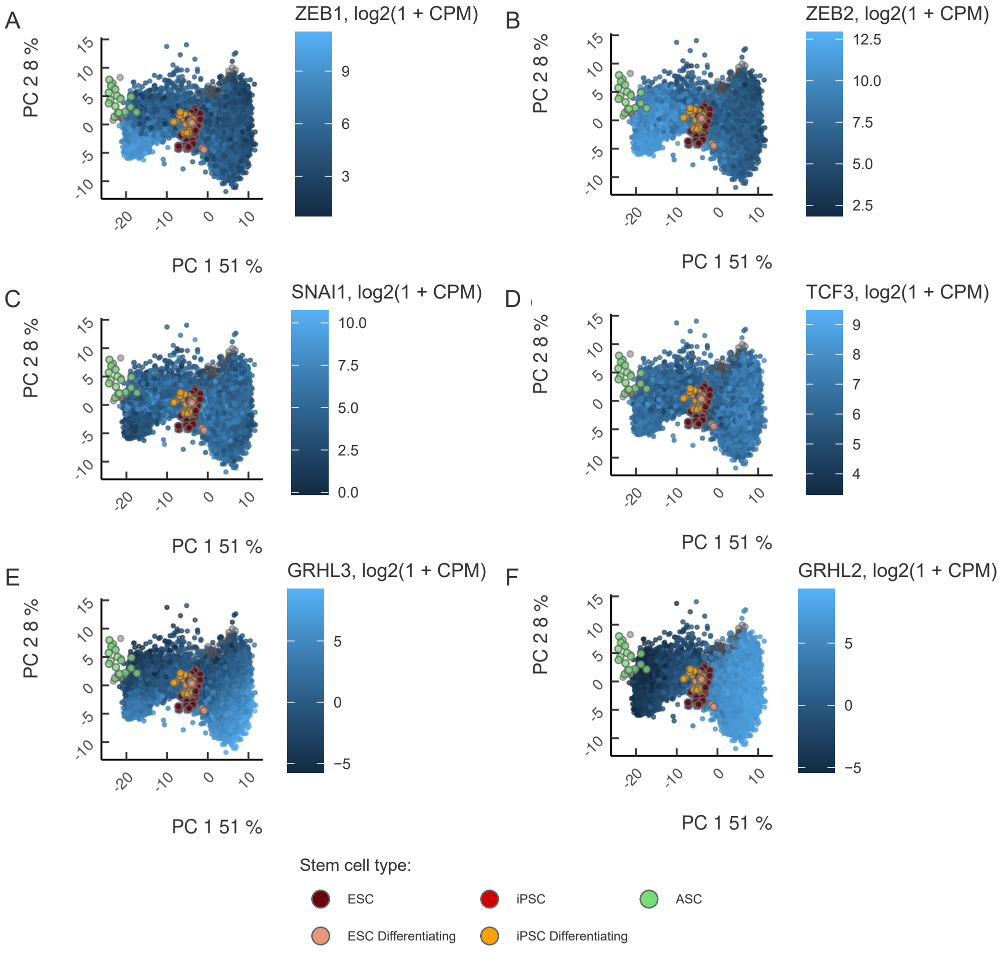


Figure S9: The gradient of expression of mesenchymal (A-D) and epithelial (E-F) genes on the PCA clustering diagram corresponding to EMT signatures (see Fig 2C). Normal tissues are shown as gray background. Stem cells are colored as in Fig 2.

| Accession | Group | Organism | $n$ | Description |
|---|---|---|---|---|
| PRJNA385016 | iPSC | Homo sapiens | 16 | Differentiation time-series (4 days, 4 replicates per day); ten passages in E8, then iDEAL feeder-free medium) |
| PRJNA383735 | ESC | Homo sapiens | 6 | Different lines (H9, HUES6, MEL1); different growth conditions (E8 and KSR media); |
| E-MTAB-5674 | ESC/PSC | Homo sapiens | 30 | Different lines (H9, SHEF6, HNES1); different growth conditions (tt2iLGo, tt2iLGo + iROCK, tt2iLGo feeder cell free, KSR/FGF); Fibroblast derived PSC (tt2iLGo) |
| E-MTAB-5114 | ESC | Homo sapiens | 12 | hESC (H9 line); undifferentiated + differentiation (CHIR, WNT3A, BMP4) |
| PRJNA695233 | ASC | Homo sapiens | 6 | Dental pulp stem cells; different age of donors (20-30 y.o., 40-50 y.o.), different growth conditions (15% serum, 60% serum) |
| PRJNA320078 | ASC | Homo sapiens | 10 | Human umbilical cord blood mononuclear cell (MNC)s-derived endothelial colony forming cells (ECFC), cultured in EGM-2 BulletKit medium (Lonza) + 10% serum |
| PRJNA531701 | ASC | Homo sapiens | 8 | Cardiac stem cells, inferior turbinate stem cells (ITSC); Cardiac stem cells cultured in DMEM/F-12 + 10% calf serum + FGF + EGF, ITSC cultured in DMEM/F-12 + 10% human plasma + FGF + EGF |

Table S1: The accession numbers and description of the public RNA-seq datasets that were used in the principal component analysis along with TCGA data; $n$ denotes the (effective) dataset size.

| Assay | Group | GEO | PMID | $\log_2 FC$ | $\log_2 E$ | Comparisons | Groups removed |
|---|---|---|---|---|---|---|---|
| Mouse GNF1M Gene Atlas | Atlas | GSE1133 | 15075390 | 0.05 | 3 | N/A | umbilical cord; E18 embryos; embryo day 10.5; embryo day 7.5; embryo day 9.5; embryo day 6.5; embryo day 8.5; fertilized egg; placenta; females with E18 embryos; testis; ovary; oocyte; thymus; b220+bcell; cd4+Tcell; cd8+Tcell |
| Mouse MOE430 Gene Atlas | Atlas | GSE10246 | 18442421 | 0.1 | 4.5 | N/A | embryonic_stem_line_V26_2_p16; umbilical_cord; stem_cells__HSC; placenta; common_myeloid_progenitor; C2C12; C3H/10T1/2; 3T3-L1; Baf3; nih_3T3;min6;mIMCD-3;RAW_264_7 |
| EB differentiation J1 | EB_diff | GSE3749 | 17394647 | 0.05 | 3 | J1_ES_0_hr_vs_J1_EB_14d | N/A |
| EB differentiation R1 | EB_diff | GSE2972 | 17394647 | 0.05 | 3 | R1_ES_0h_vs_R1_EB_14d | N/A |
| EB differentiation V6.5 | EB_diff | GSE3231 | 17394647 | 0.05 | 3 | V6.5_ES_0h_vs_V6.5_EB_14d | N/A |
| Knockdowns of pp TFs | ppTF-KDs | GSE26520 | 23462645 | 0.05 | 3 | mESC_vs_shPou5f1-mESC;mESC_vs_shNanog-mESC;mESC_vs_Esrrb-mESC;mESC_vs_shSox2-mESC;mESC_vs_shSall4-mESC | N/A |

Table S2: GEO accession numbers of the datasets that were used to assemble a set of stemness signature genes. Additional information is reported in Methods (see Signature gene sets section for details). $\log_2 FC$ denotes log-fold-change threshold (Treatment vs Control); $\log_2 E$ denotes log gene expression level threshold (Treatment + Control); Comparisons denotes the list of comparisons for differential gene expression analysis.

**SupplementaryDataFile 1:** TCGA sample subtype annotation and Stem cluster attribution

**SupplementaryDataFile 2:** Table of literature sources of manually curated stemness markers

**SupplementaryDataFile 3:** The list of stemness, proliferation and EMT/MET signature genes
